# SLAMF6 enables efficient attachment, synapse formation, and killing of HIV-1-infected CD4^+^ T cells by virus-specific CD8^+^ T cells

**DOI:** 10.1101/2025.01.20.633914

**Authors:** Blandine Monel, Pedro A. Lamothe, James Meyo, Anna P. McLean, Raymond Quinones-Alvarado, Mélanie Laporte, Julie Boucau, Bruce D. Walker, Daniel G. Kavanagh, Wilfredo F. Garcia-Beltran, Yovana Pacheco

## Abstract

Efficient recognition and elimination of HIV-1-infected CD4^+^ T cells by cytotoxic CD8^+^ T cells (CTLs) require target cell engagement and the formation of a well-organized immunological synapse. Surface proteins belonging to the SLAM family are known to be crucial for stabilizing the immunological synapse and regulating antiviral responses during lymphotropic viral infections. In the context of HIV-1, there have been reports of SLAMF6 down-regulation in HIV-1-infected CD4^+^ T cells; however, the significance of this modulation for CTL function remains unclear. In this investigation, we used CTL lines from People living with HIV (PLWH) to examine the impact of SLAMF6 blockade on three pivotal processes: (1) the formation of CD8^+^-CD4^+^ T-cell conjugates, (2) the establishment of the immunological synapse, and (3) the killing and cytokine production capacity of HIV-1-specific CTLs during HIV-1 infection. Our findings reveal that the inability to form CD8^+^-CD4^+^ T-cell conjugates following incubation with an anti-SLAMF6 blocking antibody is primarily attributable to a defect in actin ring formation at the immunological synapse. Furthermore, SLAMF6 blockade leads to a reduction in the killing efficiency of HIV-1-infected CD4^+^ T cells by HIV-1-specific CTLs, underscoring the critical role of SLAMF6 in cytolytic function. This study highlights the importance of SLAMF6 receptors in modulating cytotoxic antiviral responses, shedding light on potential avenues for manipulation and enhancement of this pathway in the context of HIV and other lymphotropic viral infections.

## INTRODUCTION

The emergence of HIV-1-specific cytotoxic CD8^+^ T lymphocytes (CTLs) during acute HIV-1 infection plays a critical role in viral control (1–3). In the chronic phase of HIV infection, CTLs are essential for controlling viral load (1–3), and viral control in a monkey model of AIDS virus infection is enhanced by interventions that augment CTL function (4)(1–3) CD8^+^ T-cell activation primarily occurs through the engagement of the T cell receptor (TCR) by cognate peptide-MHC-I complexes, leading to the elimination of HIV-infected cells. However, the precise mechanism that defines an effective functional T cell response remains incompletely understood. Research has indicated that, in addition to the TCR, several co-signaling molecules, such as CD28, LFA-1, LFA-3, PD-1, and SLAMF4, expressed on the membranes of effector and target cells, are crucial for facilitating cell-to-cell interactions required for effective recognition and killing (5–7). These molecules either induce downstream signaling, mediate adhesion, or fulfill both functions (6, 8–10). While other signaling receptors have been proposed as essential elements for the effective activation of CTLs, their specific pathways have yet to be elucidated.

SLAMF6, also known as Ly108 in mice and NTB-A in humans, belongs to the signaling lymphocyte activation molecule (SLAM) family. This family comprises co-signaling receptors expressed on the surfaces of hematopoietic cells (11). SLAMF6 engages in homophilic interactions, meaning that the SLAMF6 receptor binds to another SLAMF6 molecule, leading to the recruitment of SH-2-containing signal-transducing molecules, such as SLAM-associated protein (SAP) and Ewing’s sarcoma-activated transcript 2 (EAT-2), to its intracellular tyrosine-based switch motif (ITSM) domain. This interaction initiates a co-stimulatory signaling pathway.

SLAMF proteins have been shown to modulate the immune response, playing roles in enhancing natural killer T (NKT) cell development (12), NK cell education (13) and co-stimulating natural killer (NK) cell cytotoxicity when interacting with the influenza viral hemagglutinin (14). Disruption of the SLAMF6 signaling pathway, such as through mutations in SAP, can impede the ability of CTLs to clear Epstein-Barr virus (EBV)- infected cells, underlining the significance of the SLAMF6 pathway in mounting an effective antiviral immune response (11, 15–18). Studies in mice have demonstrated that SLAMF6 can localize to the immunological synapse and modulate T cell signaling. In SAP-deficient mice, engagement of SLAMF6 disrupts the T cell-B cell immunological synapse, resulting in reduced cytolytic function (19). In the context of cancer, a soluble SLAMF6 receptor has been shown to enhance CTL function and improve anti-melanoma activity (20), while another study highlights the necessity of SLAMF6 clustering for enhanced T cell activation (21). Conversely, when overexpressed, SLAMF6 may function as an immune checkpoint molecule, similar to PD-1, PD-L1, and CTLA-4, thereby potentially suppressing the immune system’s ability to target and attack cancer cells (22). Collectively, these studies underscore the critical role of SLAMF6 in immunological synapse formation and cytotoxicity.

In the context of HIV infection, the role of the SLAMF6 receptor and its associated signaling pathway in CTL function remains incompletely characterized, although it holds the potential to be a crucial regulatory pathway. Notably, both CD4^+^ T cells, which are the primary targets of HIV, and CTLs express high levels of SLAMF6 on their cell membranes, suggesting the possibility of interactions between SLAMF6 molecules on these two cell types (23). Furthermore, Vpu, an HIV accessory protein, has been linked to the downregulation of SLAMF6 expression in HIV-1-infected cells. However, the extent and implications of this effect on HIV-1-specific cytotoxic immune responses have yet to be thoroughly investigated (24).

In this study, we aimed to elucidate the role of SLAMF6 in HIV-1-specific CD8^+^ T cell function. Our hypothesis posited that SLAMF6 plays a pivotal role in mediating the attachment and synapse formation between effector CD8^+^ T cells and HIV-infected CD4^+^ T cells. Our results show that SLAMF6 is indeed required for virus-specific CTLs to efficiently attach to CD4^+^ T cells presenting viral peptides. Furthermore, we have demonstrated that SLAMF6 is a crucial factor in the formation of synapses and the subsequent elimination of HIV-infected CD4^+^ T cells by virus-specific CD8^+^ T cells.

## MATERIALS AND METHODS

### Human subjects

Peripheral blood was obtained by venipuncture with acid-citrate dextrose anticoagulant from People Living with HIV (PLWH) at the Massachusetts General Hospital (MGH), Boston. Written informed consent was obtained from all volunteers prior to enrollment in the study. The study was approved by the MGH Institutional Review Board (Protocol #2010-P-002463; Protocol #2010-P-002121). All subjects were on antiretroviral therapy (ART) and aviremic at the time of blood draw. Peripheral blood mononuclear cells (PBMC) from PLWH were isolated by Ficoll density gradient centrifugation within six hours of blood draw and cryopreserved in the presence of 10% DMSO in a liquid nitrogen freezer.

### Antibodies

For blockade experiments, LEAF™ Purified anti-human CD352 (NTB-A) (BioLegend, clone NT-7), anti-human CD48 functional grade purified antibody (eBioscience eBio156-4H9), LEAF™ Purified anti-human CD58 antibody (LFA-3) (BioLegend, clone: TS2/9) or mouse IgG1 K isotype control functional grade purified antibody (eBioscience clone P3.6.2.8.1) were used. For intracellular cytokine staining for flow cytometry, the following fluorochromes directly conjugated to mAbs were used: CD3-Alexa Fluor 700 (BD; clone UCHT1), CD8-Qdot 605 (Invitrogen; clone 3B5), SLAMF6-PE (BioLegend; clone NT-7), IFNγ-PE-Cy7 (BD; clone B27), CD4-APC-Cy7 (BD; clone SK3). For surface staining of human samples for flow cytometry, the following dyes, and fluorochromes directly conjugated to mAbs were used: hCD45-Alexa Fluor 700 (BioLegend; clone HI30), CD3-PerCP-Cy5.5 (BioLegend; clone UCHT1), CD4-BV785 (BioLegend; clone RPA-T4), CD8-APC-Cy7 (BioLegend; clone RPA-T8), CD45RA-PE-Cy7 (BioLegend; clone HI100), CCR7-BV421 (BD; clone 150503), CD38-BV605 (BioLegend; clone HIT2), HLA-DR-BV510 (BioLegend; clone L243), PD-1-APC (BioLegend; clone EH12.2H7), SLAMF6-PE (BioLegend; clone NT-7).

### Cell dyes

For flow cytometry CellTrace™ CFSE Cell Proliferation Kit (Life Science), CellTracker™ Violet BMQC Dye (Life Science), and PKH26 Red Fluorescent Cell dye^TM^ (Sigma Aldrich) were used. For confocal microscopy analysis CellTracker^TM^ Blue CMF2HC dye and CellTracker^TM^ Orange CMTMR dye (Invitrogen) were used.

### Peptides

Synthetic peptides were purchased from the MGH peptide core or Genscript as follows: HIV-A*02-SL9, SLYNTVATL; HIV-B*57-KF11, KAFSPEVIPMF; Influenza-A*02-GL9 GILGFVFTL; Epitope identification was facilitated by reference to the Immune Epitope Database (www.iedb.org). HIV-1 Gag 15-mer overlapping peptide pool was designed in-house and used for *ex vivo* conjugation experiments. CMV pp65 15-mer peptide pool was purchased from StemCell Technology.

### T cell culture

CTL lines specific for a given peptide were generated from cryopreserved PBMC from A*02:01 and/or B*57:01^+^ donors living with HIV. PBMCs were thawed and incubated with peptide for 10 days in R10 medium (RPMI supplemented with 10% fetal bovine serum plus Hepes buffer, penicillin, streptomycin, and L-glutamine). IL-2 (50 IU/mL) was added on day 3. Specificity for cognate peptide was tested on day 10 by intracellular cytokine staining (ICS).

### Intracellular cytokine staining

Cells were incubated with peptide (5ug/ml) for 30 min in R10, and then overnight in R10 plus 10 μg/mL brefeldin A. Cells were stained for surface markers (CD3, CD8, CD4, and SLAMF6), fixed, permeabilized, and stained for internal IFN. Flow cytometry data were acquired using a BD LSR II cytometer and were analyzed with FlowJo software (Treestar, Inc.).

Surface staining of human samples for flow cytometry: Frozen PBMC samples from 10 HIV-1-negative and 10 HIV-1-positive donors were thawed and resuspended in 1 mL of medium consisting of Advanced RPMI (Corning), 10% FBS (Avantor), 1X primocin (Invivogen), and 2 mM GlutaMAX (Life Technologies). These were transferred into a 24-well plate for a 3-hour incubation at 37°C/5% CO2 to allow rest and recovery from freeze-thaw cycle. Post-incubation, each sample was collected and passed through a 70-μm Nylon filtered FACS tube followed by a wash with 2% FBS in PBS solution. Up to 10^6^ cells of each sample were transferred into new FACS tubes, washed with 2% FBS in PBS, and resuspended in 100 μL of a master mix of 2% FBS in PBS containing 1X Live/Dead Blue (Invitrogen) and the following antibodies: hCD45-Alexa Fluor 700, CD3-PerCP-Cy5.5, CD4-BV785, CD8-APC-Cy7, CD45RA-PE-Cy7, CCR7-BV421, CD38-BV605, HLA-DR-BV510, PD-1-APC, SLAMF6-PE. The cells were then incubated for 15 minutes at 4°C. The cells were subsequently washed with 2% FBS in PBS and resuspended in 200 μL of 2% FBS in PBS before proceeding to flow cytometry, which was performed using a BD FACS Symphony (BD Biosciences) and analyzed using FlowJo v10.7.1 and GraphPad Prism 9 software.

### Flow-based conjugation assay

#### Preparation of target cells

3 days prior to the experiment, autologous CD4^+^ T cells were negatively Isolated from frozen PBMC by magnetic isolation kit (StemSep™ Human CD4^+^ T Cell Enrichment Kit, STEMCELL Technologies) and cultured in R10 supplemented with IL-2 (100 IU/ml). The day of the experiment, cells were loaded with cognate peptide at the indicated concentration for one hour at 37°C, 5% CO2. The peptide was washed off and cells were stained with Violet Dye following manufacturer’s protocol (Life Technologies).

#### Preparation of effector cells

CD8^+^ T cells were negatively isolated from epitope-specific CTL lines by magnetic separation (StemSep™ Human CD8^+^ T Cell Enrichment Kit, STEMCELL Technologies) and stained with carboxyfluorescein succinimidyl ester (CFSE).

#### Co-incubation

CD8^+^ and CD4^+^ T cells were co-cultured at a ratio 1:1 in a flat bottom 96-well plate; the cells were centrifuged for 3 minutes at 25G and then incubated for 15 min in a 37°C, 5% CO2 incubator. Then, the plate was placed on ice, spun down at 4°C for 5 minutes at 750G. After, the supernatant was aspirated and cells were stained with a viability dye 7-AAD (BD Pharmingen™), incubated for 10 min at 4°C, washed, and gently resuspended in PBS-4% Paraformaldehyde (PFA) for fixation on ice. Samples were analyzed by flow cytometry using a BD 5 Laser LSR2 Fortessa instrument. Flow cytometry data were analyzed by FlowJo software (Treestar, Inc). The rate of antigen-specific conjugation was determined by the frequency of double positive events consisting of both green effector cells and violet loaded target cells.

### Confocal microscopy and analysis

CD4^+^ T cells were stained with 5uM CellTracker^TM^ Blue CMF_2_HC dye and CD8^+^ T cells with 5μM CellTracker^TM^ Orange CMTMR dye according to the manufacturer’s instructions. CD4^+^ T cells were then incubated with or without cognate peptides (5 ug/ml) for 1h at 37°C. The peptide was washed off, and LEAF™ Purified anti-human CD352 antibody or isotype IgG control was added. CD4^+^ T cells and CD8^+^ cells were then mixed at a 1:1 ratio and were loaded onto polylysine-coated coverslips (1x10^6^ cells in 160 μL) in 24 well plates. After 15 min at 37°C, cells were fixed with 4% PFA for 15 min at room temperature, permeabilized with triton 0.5% for 15 minutes at room temperature, blocked with 3% BSA in PBS for 1 hour at room temperature, and stained with AlexaFluor 488 Phalloidin (Molecular Probes) overnight at 4°C. The coverslips were then mounted on slides using ProLong Gold Antifade Reagent (Life Technology). Images were acquired with a Zeiss LSM 510 confocal microscope equipped with argon (488 nm) and Diode (405 nm, 561 nm) lasers, 100 X oil immersion objectives, and ZEN software. At least 10 contacts between CD4^+^ T cells and CD8^+^ T cells were analyzed per condition with Z-stacks spaced by 0.33μm. The 3-D images were then reconstituted using the IMARIS software and surfaces were created for each channel after background deduction.

#### Actin polarization quantification

3D reconstituted images of synapses were analyzed with Imaris software. The fluorescent signal of the Phalloidin was quantified at the site of contact between the effector and the target cell and compared to the Phalloidin signal on the rest of the CD8 T cell. Ten immunological synapses were analyzed per condition and the mean of the ratio of actin at synapse to actin in the rest of the CD8^+^ cell was calculated for each coupling.

#### Actin ring quantification

Confocal Images were analyzed by Imaris Software, *en face* views from each duplex were taken and the blue pixels (target cells) were removed to allow an unobstructed view of the actin stain at the interface. Three blinded reviewers were asked to score the actin network at the synapse, where 10 was the maximum given to a perfect actin ring and 1 was the lowest score corresponding to the absolute absence of actin rings. A total of 100 duplex images formed under different conditions were randomized and quantified for the actin ring formation.

### HIV-1 infection

Phytohemagglutinin-activated CD4^+^ T cells were harvested and incubated at 1 × 10^6^ cells/mL in 200 μL of R10 in the presence of HIV-1 strain NL4.3 for 5 hours. Cells were then washed and plated at a concentration of 1× 10^6^ cells/ml in a 6-well plate for 48 hours.

### Chromium-release assay

#### Targets

CD4^+^ T cell isolated from PBMCs were loaded with cognate peptide at the indicated concentration and incubated with ^51^Cr at a concentration of 250 μCi /ml for one hour in a 37°C 5% CO2 incubator. ^51^Cr-Labeled cells were washed 3 times in R10 and resuspended at a concentration of 1 million cells/ml. Target cells were plated in a flat bottom 96-well plate and incubated with an isotype control or anti-SLAMF6 blocking antibodies for 15 min. Then, effector CD8^+^ T cells isolated from epitope-specific CTL lines were added at the right concentration in order to have an effector-target ratio of 1:1. Target cells treated with 5% Triton solution were used to measure maximum chromium release and target CD4^+^ T cells incubated without effectors were used for measuring the spontaneous ^51^Cr release. Cells, whether loaded with peptide or infected, were plated in the absence of effector cells to determine the level of spontaneous release (see S**upplemental Figure 4**). The supernatant was collected after 4 hours of incubation at 37°C 5%CO2. We used a Perkin Elmer TopCount NXT Microplate Scintillation & Luminescence Counter to measure radioactivity present in the supernatant. Quantification of specific killing was calculated by specific killing =100* (sample release – spontaneous release) /(maximum release-spontaneous release).

### Statistics

Quantitative variables were compared by two-tailed Wilcoxon matched-pairs signed rank test when quantitative variables were paired or Mann-Whitney test when comparisons were made between independent quantitative variables. Correlations were calculated using the Spearman correlation test. All calculations were performed using GraphPad software version 6 and a difference or a correlation was considered significant when p < 0.05. For every quantitative variable shown the error bar indicated the SEM (Standard Error of the Mean).

## RESULTS

### Epitope-specific attachment and conjugation of CTL lines and CD4^+^ T cell targets

In order to assess the ability of CD8^+^ T cells to recognize and attach to viral peptides-presenting CD4^+^ T cells, we developed a flow-based conjugation assay with cells from people living with HIV (PLWH) that measures the rate of CD8^+^-CD4^+^ T cell doublets between CTL lines specific for HLA-A*02:01-Flu-GL9 or HLA-B*57:01-HIV-KF11 and autologous CD4^+^ T cell targets presenting MHC-class I epitopes (**Figure 1A**).

**Figure 1:**
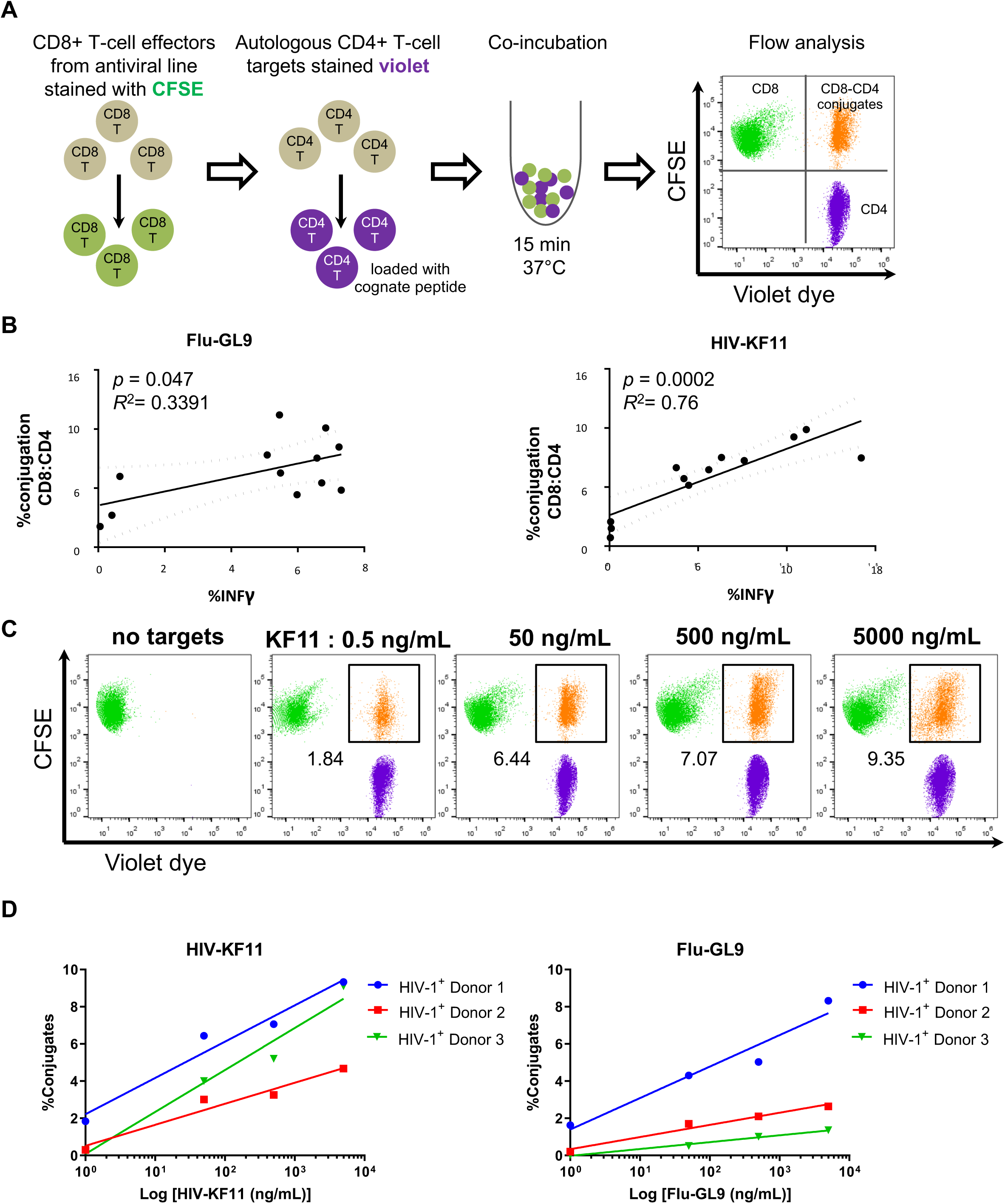
Conjugation of virus-specific CD8^+^ CTLs to CD4^+^ T-cell targets loaded with cognate peptide can be accurately measured via flow cytometry and predict CTL functionality. (**A**) Experimental two-color design for quantifying antigen-specific CD8^+^ and CD4^+^ T-cell Interactions is shown. In this experiment, CTLs were labeled with green fluorescence (CFSE) and autologous CD4^+^ T cells were loaded with varying doses of cognate peptide and stained with a violet dye. Effector and target cells were mixed at a 1:1 ratio of CD8^+^ to CD4^+^ T cells and co-cultured at 37°C for 15 minutes. Subsequently, the cells were fixed and subjected to flow cytometry analysis. The rate of antigen-specific conjugations was determined by calculating the frequency of double-positive cells, where green effector cells overlapped with violet target cells, located in the upper right quadrant of the cytometry plot. (**B**) We investigated the relationship between the frequency of CD8^+^-CD4^+^ T cell conjugate formation occurring within a 15-minute timeframe and subsequent cytokine production by CD8^+^ CTLs at six hours of co-culture. This was done for HIV-1 KF11 peptide (HIV-KF11)-specific CTLs (left) and influenza GL9 peptide (Flu-GL9)-specific CTLs (right) at varying concentration of cognate peptide loaded onto autologous CD4^+^ T-cell targets (*n* = 12 for each). The X-axis represents the frequency of specific conjugate formation observed after 15 minutes, while the Y-axis illustrates the frequency of IFN-γ^+^ CTLs after 6 hours of co-culture. The graphs presented here serve as representatives from a total of three independent experiments, and the correlation between these variables was assessed using a simple linear regression (*p* values and *R*^2^ shown on graphs). (**C**) CD4^+^ T cell targets were loaded with the HIV-KF11 epitope at the indicated concentrations and then washed to remove excess peptide. HIV-KF11-specific CD8^+^ CTLs were incubated with autologous peptide-pulsed CD4^+^ T cell targets for 15 minutes. Each flow plot represents a different concentration of the loaded peptide, and the conjugate formation is depicted in orange (identified as the double-positive green and violet population) and the conjugate frequency is indicated for each plot. This experiment is representative of three independent experiments. (**D**) The effect of cognate peptide concentration on conjugates formation was measured by co-incubating HIV-KF11-specific (left) or Flu-GL9-specific (right) CTLs with autologous CD4^+^ T-cell targets loaded with different concentration of HIV-KF11 or Flu-GL9 peptide. A simple linear regression between the log-transformed peptide concentration (X-*axis)* and percent of conjugates (Y-*axis*) was assessed, with results as follows: KF11 Donor 1 (*R*^2^ = 0.96, *p* = 0.02), KF11 Donor 2 (*R*^2^ = 0.95, *p* = 0.02), KF11 Donor 3 (*R*^2^ = 0.96, *p* = 0.02), GL9 Donor 1 (*R*^2^ = 0.94, *p* = 0.03), GL9 Donor 2 (*R*^2^ = 0.97, *p* = 0.01), and GL9 Donor 3 (*R*^2^ = 0.99, *p* = 0.006).

To validate the sensitivity of the assay, we analyzed in parallel the formation of conjugates after 15 minutes and the frequency of CD8^+^ T cells producing IFN-γ after 6 hours of stimulation. We observed that the rate of CD8^+^-CD4^+^ T cell conjugate formation after 15 minutes of co-culture predicted the rate of IFN-γ production by CD8^+^ T effector cells 6 hours after the stimulation (*p* =0.0002 and R^2^=0.76 for HIV-KF11; *p* = 0.05 and R^2^=0.34 for Flu-GL9) (**Figure 1B**). There was a statistically significant correlation between conjugate formation and IFN-γ production for the different tested peptides from HIV-1 and influenza. This correlation was stronger for HIV-KF11-specific CTLs than for Flu-GL9-specific CTLs. In addition, conjugate formation was proportional to the peptide concentration used to pulse the target cells (**Figure 1C, 1D**) with the following linear regression results: KF11 Donor 1 (*R*^2^ = 0.96, *p* = 0.02), KF11 Donor 2 (*R*^2^ = 0.95, *p* = 0.02), KF11 Donor 3 (*R*^2^ = 0.96, *p* = 0.02), GL9 Donor 1 (*R*^2^ = 0.94, *p* = 0.03), GL9 Donor 2 (*R*^2^ = 0.97, *p* = 0.01), and GL9 Donor 3 (*R*^2^ = 0.99, *p* = 0.006).

Having validated our experimental approach, we proceeded to assess and differentiate epitope-specific and non-specific conjugation events. We set up an assay similar to the cold-target competition assay used to measure cytotoxicity (25). In our setup, violet-labeled non-pulsed CD4^+^ T cells (“cold targets”), red-labeled CD4^+^ T cells pulsed with HIV-KF11 (“warm targets”), and green-labeled HIV-KF11-specific CD8^+^ T-cell effectors were co-incubated for 15 minutes, and conjugate formation was assessed (**Figure 2A**). Supplemental Figure 1 shows the gating strategy used for all the conjugation assays. We demonstrated that blockade with anti-LFA-3 (also known as CD58) antibody, a known adhesion receptor, dramatically reduced peptide-specific conjugation (**Figure 2B**). Importantly, LFA-3 was used as a positive control, as it is an adhesion molecule necessary to create effective contact between T lymphocytes and antigen-presenting cells (26, 27). Twenty CTL specific lines from 10 HIV-positive patients were tested, and for each of them, we observed that LFA-3 blockade significantly reduced peptide-specific conjugation compared to the isotype control antibody. This finding aligns with the previously described role of LFA-3 as an adhesion molecule (26, 27). Furthermore, although the blockade of LFA-3 resulted in a decrease in the rate of non-specific conjugation, the overall rate of non-specific conjugation was minimal. This observation further affirms the validity of our experimental approach.

**Figure 2:**
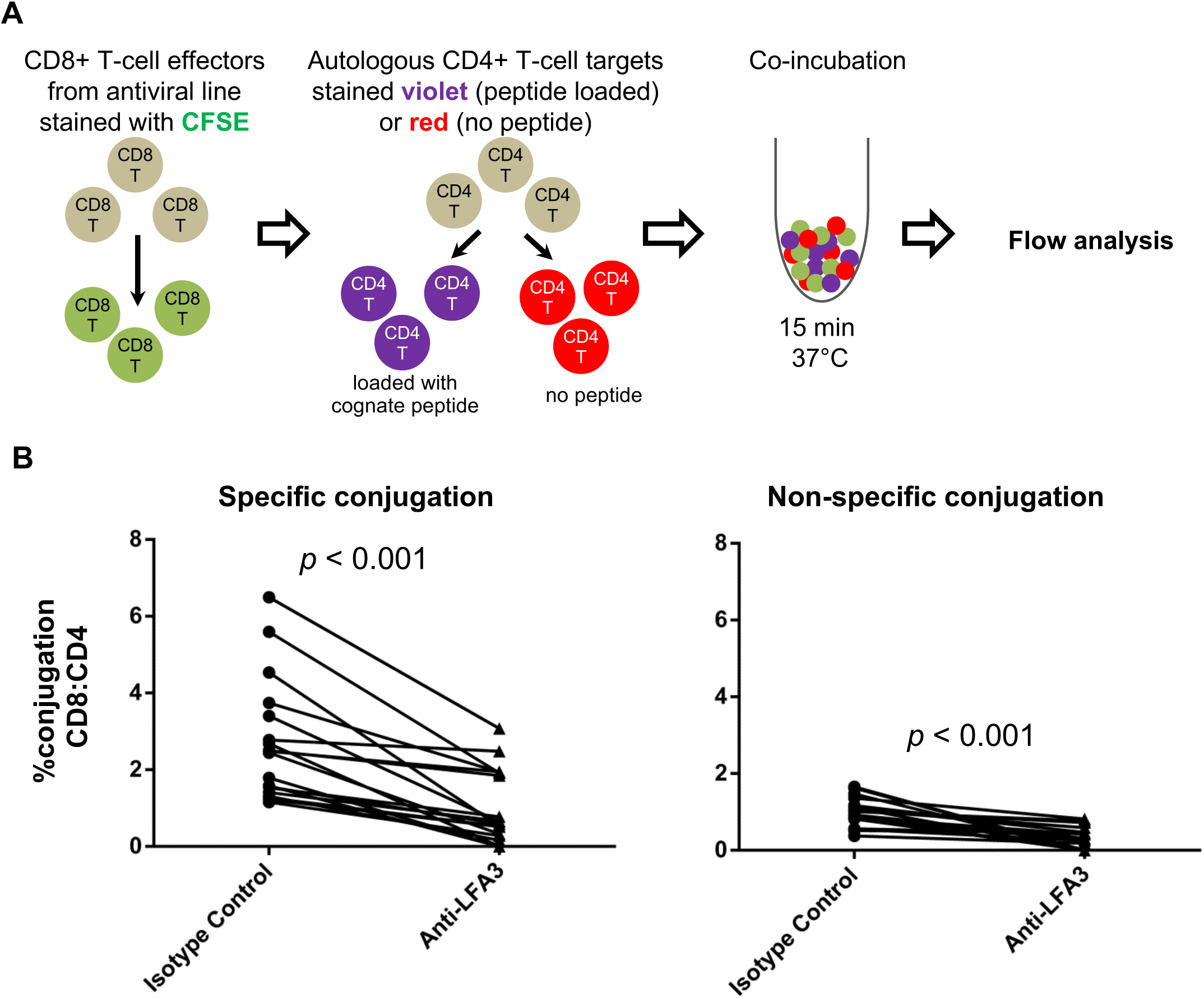
Blockade of LFA-3 inhibits both antigen-specific and non-specific attachment and conjugation of CD8^+^ CTLs to autologous CD4^+^ T cells. (**A**) We developed a three-colors experimental design for measuring antigen specific and non-specific attachment and conjugation of CD8^+^ CTLs to autologous CD4^+^ T cells. CTLs were labeled with green fluorescence (CFSE) and autologous CD4^+^ T cells isolated from donor PBMCs were divided into two samples: peptide-pulsed targets cells stained in violet and non-peptide-pulsed target cells stained in red. Peptide-pulsed target cells and non-pulsed target cells were co-incubated at a ratio 1:1:1 for 15 minutes at 37°C. Cells were fixed and analyzed by flow cytometry. Within the CFSE+ cells (i.e., CD8^+^ CTLs) the rate of antigen-specific conjugation is determined by the frequency of violet-positive conjugates and the rate of non-specific conjugation is determined by the frequency of red-positive conjugates. (**B**) CD8^+^-CD4^+^ conjugate formation sensitive to LFA-3 blockade was explored. CD8^+^ T cells were co-cultured with CD4^+^ T target cells for 15 minutes. Conjugate formation was analyzed by flow cytometry, gating on cells that were double positive for CD8 and CD4. Two conditions were assessed: (*i*) Percent of conjugation of CD8^+^ T cells (CFSE^+^) and CD4^+^ T cells loaded with cognate peptide (Violet+) (left panel), and (*ii*) Percent of conjugation of CD8^+^ T cells (CFSE^+^) and CD4^+^ T cells not loaded with cognate peptide (Red^+^) (right panel). The analysis was performed using either an isotype control antibody or a blocking antibody against LFA-3 (CD58). Co-cultures were analyzed to determine the percentage of specific and non-specific conjugates formed under these conditions. Data represents the conjugation rate from 20 independent experiments (*n* = 20).

### Role of SLAMF6 in effector-target conjugate formation

Prior studies have demonstrated that HIV-1 Vpu downmodulates SLAMF6 on HIV-1-infected CD4^+^ T cells (24, 28). We tested this by infecting *in vitro* activated primary CD4^+^ T cells from healthy donors with GFP-expressing HIV-1 and measuring SLAMF6 expression 48 h post-infection. We observed that SLAMF6 is downmodulated on CD4^lo^ GFP^+^ (i.e. infected) cells when compared to CD4^hi^ GFP– (i.e. non-infected) cells, thus confirming previous findings (**Supplemental figure 2**). In light of this finding and previously published data demonstrating the importance of SLAMF6 signaling in immunological synapse formation (17), we hypothesized that SLAMF6 plays a role in virus-specific CD8^+^ T-cell attachment to target cells presenting viral peptides. To assess the role of SLAMF6 in CD8^+^ T cell attachment to target cells we carried out the flow cytometry-based conjugation assay in the presence of anti-SLAMF6 or an isotype control antibody and determined the rate of specific conjugation of virus-specific CTL lines specific for HIV-SL9, HIV-KF11, or Flu-GL9 to autologous CD4^+^ T cell targets pulsed with cognate viral peptide. Therefore, twenty CTL lines were tested, and we observed that SLAMF6 blockade reduced the rate of peptide-specific conjugation (average reduction of 31.7 % ± 5.1 %); *p* < 0.001; **Figure 3A**). However, SLAMF6 blockade did not significantly affect non-specific conjugation (**Figure 3B**). This was not due to differences in SLAMF6 expression across CTL lines since SLAMF6 was found to be highly expressed and at comparable levels across all CTL lines (MFI = 2455 ± 93) and *in vitro* cultured CD4^+^ T cells (MFI = 2301 ± 82) (**Figure 3C** and **3D**). Overall, these results suggest that SLAMF6 plays an important role in the adhesion of virus-specific CD8^+^ T cells to viral peptide-presenting CD4^+^ T cells.

**Figure 3:**
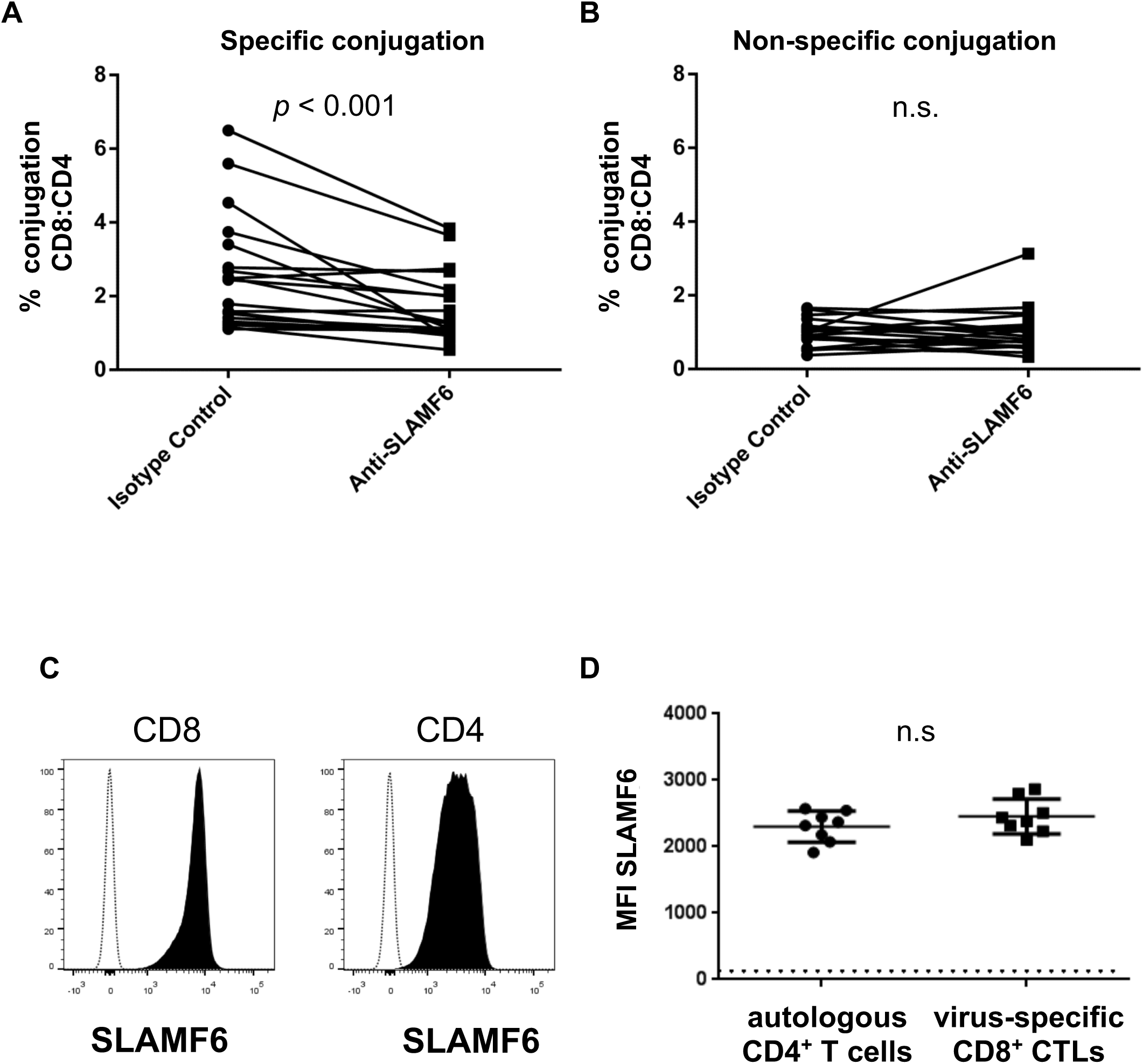
SLAMF6 is required for efficient formation of primary human antiviral CD8^+^-CD4^+^ T cell conjugates. (**A**-**B**) The conjugation assay was performed with primary T cell lines from a cohort of HIV1-positive subjects (*n* = 10). Multiple CTL lines were generated for each donor and were specific for any of the following viral epitopes: HIV-SL9; HIV-KF11; Flu-GL9. Compiled results from a total of 20 CTL lines are shown. The rate of specific conjugation in the presence of SLAMF6 blockade or isotype control is shown in (**A)** and the rate of non-specific conjugation is shown in (**B**) The effect of the blockade was compared to the IgG isotype control by a two-tailed Wilcoxon matched-pairs signed rank test; *p* values are shown on graph, with n.s. denoting non-significant (*p* > 0.05). (**C**) Representative flow cytometry histograms measuring SLAMF6 expression on T cells used for the conjugation assay is shown for virus-specific CD8^+^ CTLs (left) and autologous primary CD4^+^ T cells isolated from PBMC and maintained for 3 days in medium containing IL-2 (50 U/ml) (right). The black-filled histogram represents SLAMF6 staining and the dotted histogram represents the FMO (fluorescence minus one). (**D**) Combined results of SLAMF6 expression on CTLs and IL-2-cultured CD4^+^ T cells from 10 different HIV-1-positive subjects are shown. Two-tailed Wilcoxon matched-pairs signed rank test showed no statistically significant differences (n.s.).

The sensitivity of CTL lines to the SLAMF6 blockade varied depending on HLA class I specificity. We observed that CTL lines that recognized the HIV-KF11 restricted by HLA-B*57:01 haplotype were less sensitive to the SLAMF6 blockade than CTL lines recognizing HIV-SL9 and Flu-GL9 restricted by HLA-A*02:01 haplotype (**Figure 4A**). For HLA-A*02:01-restricted CTL lines, the SLAMF6 blockade reduced the frequency of CD8^+^-CD4^+^ T cell conjugates by about 39.9 % ± 16.8 % relative to the isotype control, whereas HLA-B*57:01-restricted CTL lines decreased conjugate formation in the presence of SLAMF6 blockade by only 16.7 % ± 6.3%. In contrast, sensitivity towards LFA-3 blockade was comparable for both specificities (**Figure 4B**).

**Figure 4:**
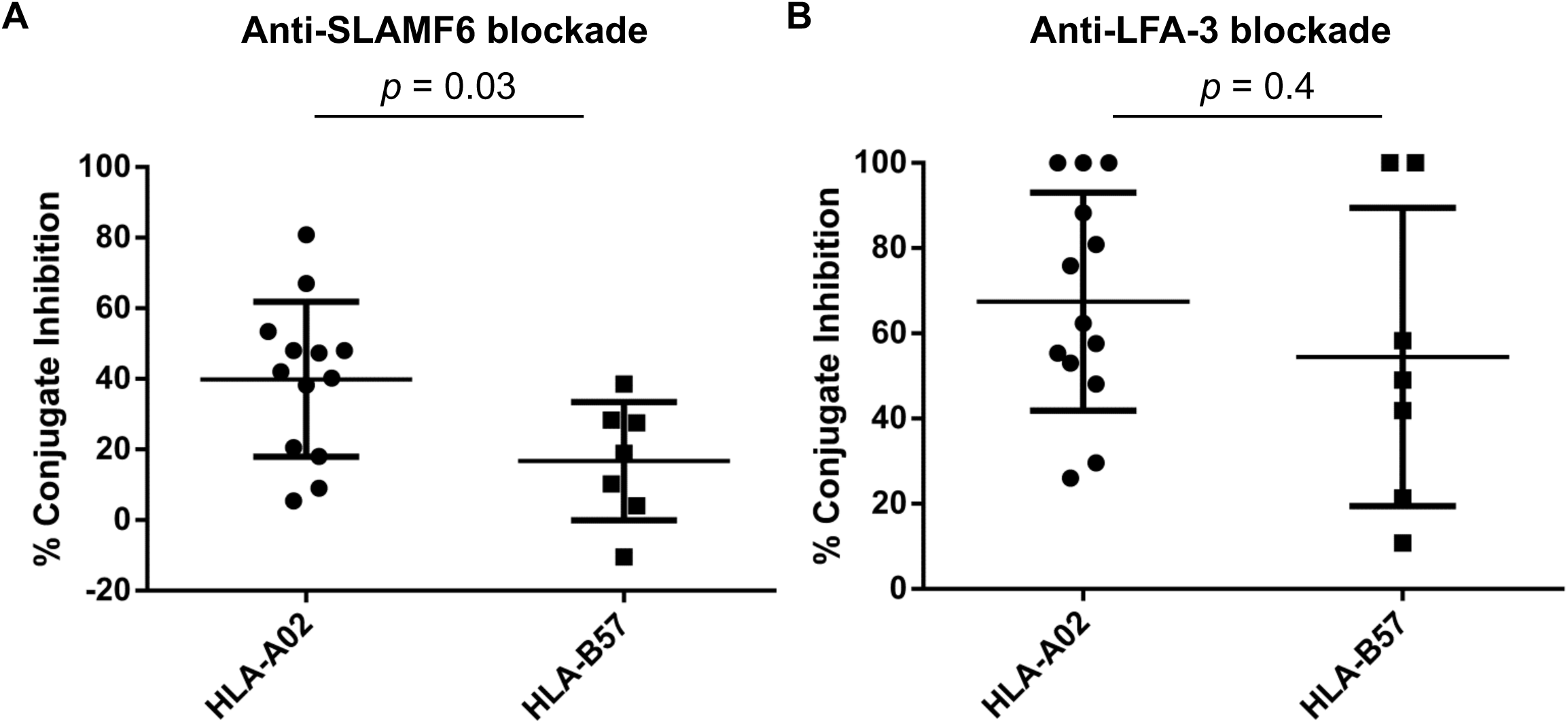
HLA-B57:01-restricted CTL lines are less sensitive to SLAMF6 blockade than HLA-A*02:01-restricted lines. For each CTL line studied, the “percent conjugate inhibition” was calculated as: 100% - (Percent specific conjugation in the presence of the blockade / Percent of specific conjugation in the presence of the IgG Isotype control). The mean “specific-conjugate inhibition” in the presence of blockade of (**A**) SLAMF6 or (**B**) LFA-3 was assessed for HLA-B57:01-restricted CTL lines (*n* = 7) compared to HLA-A*02:01-restricted CTL lines (*n* = 13) and compared by the Mann-Whitney test. The *p*-value is indicated for each graph.

To further investigate the role of SLAMF6 in antiviral human T cell responses in a more physiological model, we utilized bulk PBMCs *ex vivo*. For this purpose, we did not limit the stimulation to a single cognate peptide but used a mixture of overlapping peptides from HIV-1 Gag protein and a mixture of peptides from CMV pp65 protein. Of note, we chose CMV peptides, considering the high prevalence of this infection in the general population (greater than 45%) (29).

Firstly, we measured SLAMF6 expression levels on peripheral blood CD4+ and CD8+ T cells from HIV-1-negative (n = 10) and HIV-1-positive (n = 10) human donor PBMCs *ex vivo* (**Supplemental Figure 3A**). Due to previous studies demonstrating differential SLAMF6 expression across different T cell subsets (30), we subclassified CD4+ and CD8+ T-cell subsets as naïve (CD45RA+CCR7+) or non-naïve (CD45RA– or CCR7–), which includes memory and effector T cells, and among non-naïve cells, we identified activated (HLA-DR+CD38+) T cells and exhausted (PD-1+) T cells (**Supplemental Figure 3B**). We found that among HIV-1-negative and HIV-1-positive donors, naïve CD4+ and CD8+ T cells expressed SLAMF6 at similar levels (**Figure 5A**). However, non-naïve CD4+ T cells and CD8+ T cells had significantly higher levels (approximately 2-fold higher relative to naïve), with non-naïve CD8+ T cells having significantly higher levels of SLAMF6 than non-naïve CD4+ T cells in both donor cohorts (**Figure 5A**). Furthermore, activated and exhausted CD8+ T cells exhibited similar SLAMF6 expression to non-naïve CD8+ T cells. In contrast, activated and exhausted CD4+ T cells expressed significantly higher levels of SLAMF6 than bulk non-naïve CD4+ T cells, with SLAMF6 levels more similar to those seen among non-naïve CD8+ T cells (**Figure 5A**). This suggests that non-naïve, activated, and exhausted CD8+ T cells constitutively express similarly high levels of SLAMF6 at baseline, CD4+ T cells further increase SLAMF6 expression levels upon activation and subsequent exhaustion. This is congruent with our prior data demonstrating that *in vitro* activation and expansion of CD4+ and CD8+ T cells with IL-2 results in similarly high expression levels of SLAMF6 ( **Figure 3C, 3D**). Regarding HIV-1 donor status, there was a trend for decreased SLAMF6 expression among naïve, non-naïve and activated CD8^+^ T cells and increased SLAMF6 expression on exhausted CD4+ T cells among HIV-1-positive donors, but this was not statistically significant (**Figure 5A**). We did, however, find significant increases of activated CD4+ and CD8+ T cells frequencies in HIV-1-positive donors (**Supplemental Figure 3C**), which is characteristic of HIV-1 infection (30). Overall, SLAMF6 is highly expressed among all CD4+ and CD8+ T-cell subsets, with even higher expression among non-naïve, activated, and exhausted T cells, and no significant differences between HIV-1-negative and HIV-1-positive individuals.

**Figure 5:**
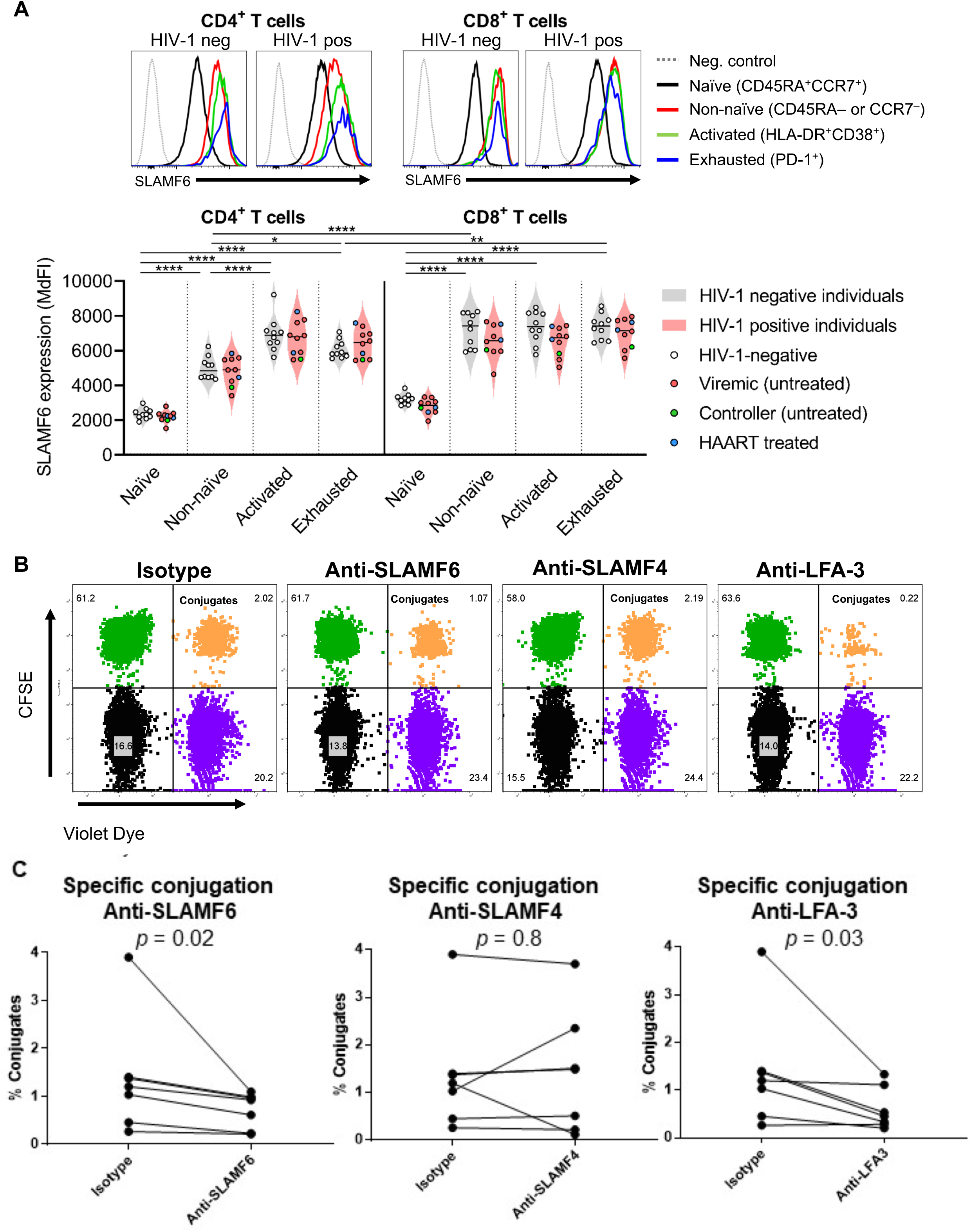
SLAMF6 is highly but differentially expressed on CD8^+^ and CD4^+^ T-cell subsets and is required for efficient CD8^+^-CD4^+^ T cell conjugate formation directly e*x vivo*. (**A**) SLAMF6 expression, measured as median fluorescence intensity (MdFI), was assessed on different CD8^+^ T and CD4^+^ T-cell subsets in PBMCs from HIV-1-positive (*n* = 10) and HIV-1-negative (*n* = 10) donors *ex vivo* via flow cytometry; representative histograms of SLAMF6 expression on different T-cell subsets is shown, with aggregate data plotted below as violin plots. One-way ANOVA with correction for multiple comparisons was performed; statistical significance is denoted as follows: * *p* < 0.05, ** *p* < 0.01, **** *p* < 0.0001. (**B**-**C**). Conjugation assay on CD8^+^ T cells from *ex vivo* PBMCs and autologous CD4^+^ T target cells in the presence of SLAMF6, SLAMF4 or LFA-3 blocking antibodies, or isotype IgG control. The peptides used in this assay were pools of overlapping synthetic peptides representing the HIV-1 Gag protein or the CMV pp65 protein. Specific conjugation is shown in the upper left quadrant. All subjects studied were HIV-1 and CMV co-infected. The flow plots in **B** are representative of six additional conjugation assays performed on *ex vivo* T cells. The rate of specific conjugation in the presence of a SLAMF6, SLAMF4 and LFA-3 blockade was assessed and shown in **C** (*n* = 6). The effects of the blockade were compared to an IgG isotype control using a two-tailed Wilcoxon matched-pairs signed rank test with *p-*values shown on graphs.

Having established the dynamics of SLAMF6 expression among peripheral blood CD8+ and CD4+ T cells, we obtained PBMCs from a cohort of HIV-1-positive patients who were also seropositive for CMV. We then conducted a conjugation assay using target CD4+ T cells loaded with overlapping synthetic peptides from HIV-1 Gag or CMV pp65 protein. The effect of LFA-3, SLAMF4, and SLAMF6 blockade on ex vivo T cells was assessed in three different subjects co-infected with HIV-1 and CMV.

As shown for one representative patient (Figure 5B) and in the combined results (Figure 5C), the LFA-3 blockade abrogated specific conjugation for HIV-1 Gag-specific *ex-vivo* CD8+ T cells confirming the role of LFA-3 in cell-cell interaction (26, 27), while SLAMF4 blockade had no significant effect, indicating that the SLAMF4 receptor does not participate in the formation of conjugates and showing that the effect of SLAMF6 is specific. However, our combined results demonstrated that SLAMF6 blockade decreased specific conjugation between virus-specific CD8+ T cells and viral peptide-presenting CD4+ T cells by approximately 46.17 ± 4.7% compared to the isotype control (Figure 5C).

Overall, these findings indicate that SLAMF6 plays an important role in the formation of stable conjugates between effector CD8^+^ T cells and CD4^+^ T cell targets, both in virus-specific CTL lines and *ex-vivo* lymphocytes.

### Effect of SLAMF6 blockade on immunological synapses formation

Effective killing of target cells by CD8^+^ T cells requires peptide-MHC recognition by T cell receptor (TCR), followed by the establishment of a cytolytic immunological synapse: which involves polarization of actin towards the point of contact and rearrangement of this actin to form a connecting membrane ring with a central clearance through which cytolytic granules are released (31). Given the previously described role of co-signaling molecules in this process, we investigated the role of SLAMF6 in immunological synapse formation between CD8^+^-CD4^+^ T cell conjugates using high-resolution confocal microscopy. To this end, we used HIV-1-specific CTL lines and HIV-peptide-pulsed autologous CD4^+^ T cell targets and imaged immunological synapses formed in the presence of anti-SLAMF6 or isotype control antibody.

Although SLAMF6 blockade decreased the number HLA-B*57:01-CTL conjugates, because the HLA-B*57:01-restricted CTL lines were less sensitive to the SLAMF6 blockade we were able to locate under the microscope several CD8-CD4 contacts even after blockade with SLAMF6. For this reason, we used B*57:01-restricted CTL lines specific for KF11 peptide to analyze the actin cytoskeleton reorganization with or without SLAMF6 blockade. Indeed, we showed an actin polarization with capping of the actin cytoskeleton towards the zone of contact between autologous CD4+T cells pulsed with KF11 peptide and the CTL cells. We observed that blockade of SLAMF6 decreased actin polarization in CD8^+^ T cells towards the contact site with CD4^+^ T cells (**Figure 6A and 6B**). In addition, central clearance of actin at the immunological synapse was blunted in the presence of SLAMF6 blockade, which led to the absence of the actin ring (**Figure 6C and 6D**). Altogether, these results show that SLAMF6 blockade disrupts actin polarization and cytolytic immunological synapse formation in virus-specific CD8^+^ T cells contacting viral peptide-presenting CD4^+^T cells.

**Figure 6:**
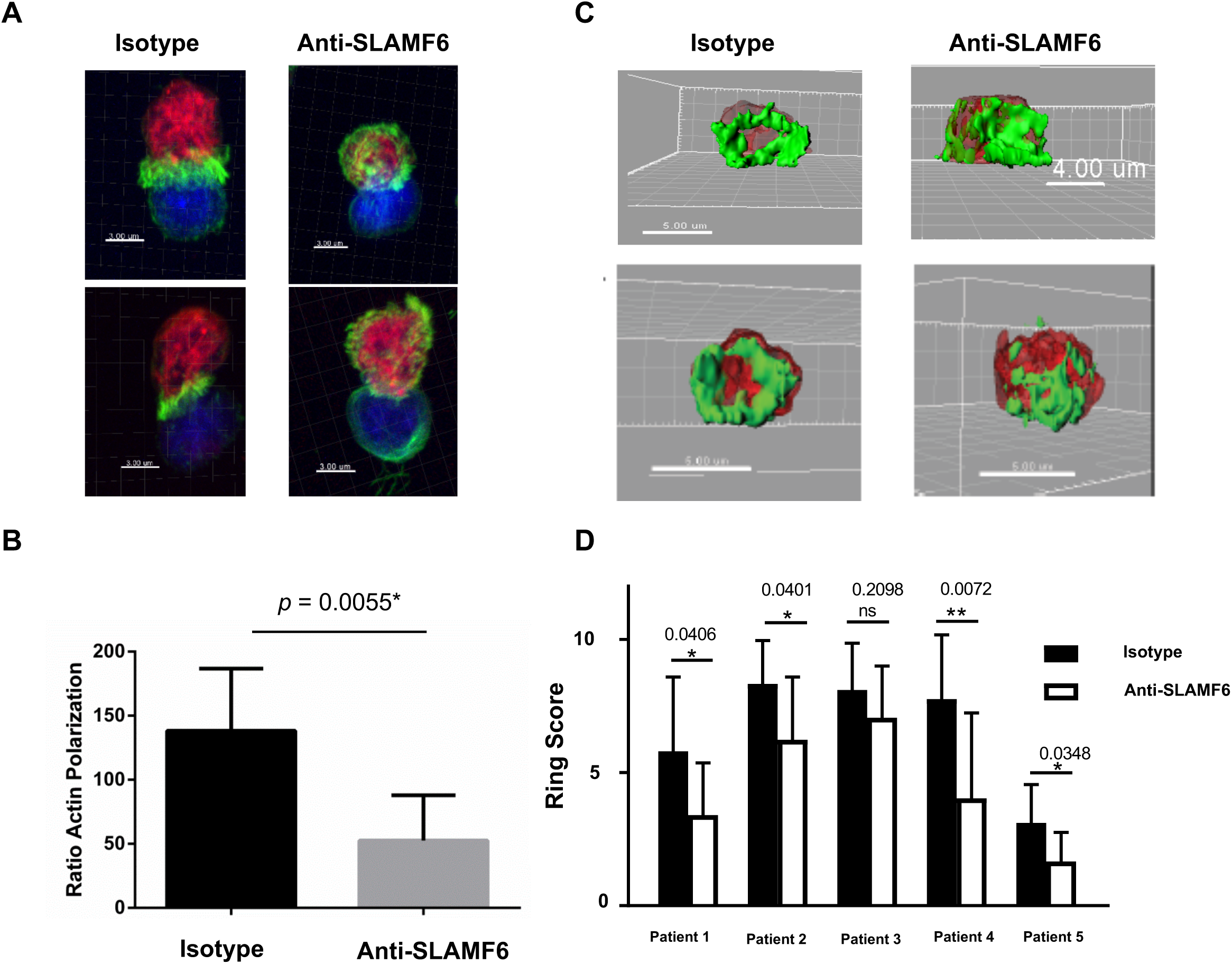
SLAMF6 is necessary for immunological synapse formation between CD8^+^ T effector cells and CD4^+^ T target cells. (**A**) HIV-KF11-specific CD8^+^ T cells (red) were co-cultured for fifteen minutes with autologous antigen-loaded CD4^+^ T cell targets (blue) in the absence or presence of a SLAMF6 blockade, stained for actin localization with phalloidin (green), and analyzed by confocal microscopy. Four different images with a magnification of X100 are shown, representative of 100 different pictures taken. (**B**) Antigen-specific actin polarization in CD8^+^-CD4^+^ T cell contacts is observed in the presence of the SLAMF6 blockade (right) and isotype control (left). The ratio of actin polarization in the presence or absence of the SLAMF6 blockade was compared using the Mann-Whitney test; the *p*-value is indicated in the graph. (**C**) The 3D plots show *en face* views of the CD4^+^-CD8^+^ T cell contacts with a magnification of 100X. The blue pixels were removed to allow an unobstructed view of the synapse. Four representative images are shown, both in the presence of the isotype control (left) and the SLAMF6 blockade (right). (**D**) The ring formation was scored for all 100 images analyzed from 5 independent experiments, and SLAMF6 vs. isotype control were compared for each experiment and analyzed by the Mann-Whitney test; the *p*-value is indicated in the graph.

### SLAMF6 blockade decreases CTL cytotoxicity

To assess the functional consequences of decreased conjugate formation by SLAMF6 blockade, we measured the cytotoxic activity of CTL lines specific for HIV-KF11 and HIV-SL9 by chromium-51 (^51^Cr) release assay in the presence of anti-SLAMF6 or isotype control antibodies. We found specific killing of CTL lines against peptide loaded and against HIV-infected target cells but no killing against targets in the absence of peptide or infection. (**Supplemental Figure 4**). We observed a decrease in specific killing in the presence of anti-SLAMF6 antibody for CTL lines restricted by both HLA-A*02:01 (41.8%±17.95 reduction in the presence of SLAMF6 antibody) and HLA-B*57:01 (51.7.8%± 20.32 reduction in the presence of SLAMF6 antibody) alleles (**Figure 7A**). To further investigate the role of SLAMF6 in a more biologically-relevant model, we compared the killing of HIV-infected CD4^+^ T cells by autologous specific CTL lines in presence of an isotype control or an anti-SLAMF6 blockade antibody. We showed that specific killing was reduced from 11.18 %±8.45 to 4.2%±1.92 (35.59%±6.41 rate of reduction) in the presence of SLAMF6 blockade antibody (**Figure 7B**). These data show that SLAMF6 plays a role in the cytolytic function of CD8^+^ T cells against HIV-infected CD4^+^ T target cells by participating to the establishment of functional immune synapses.

**Figure 7:**
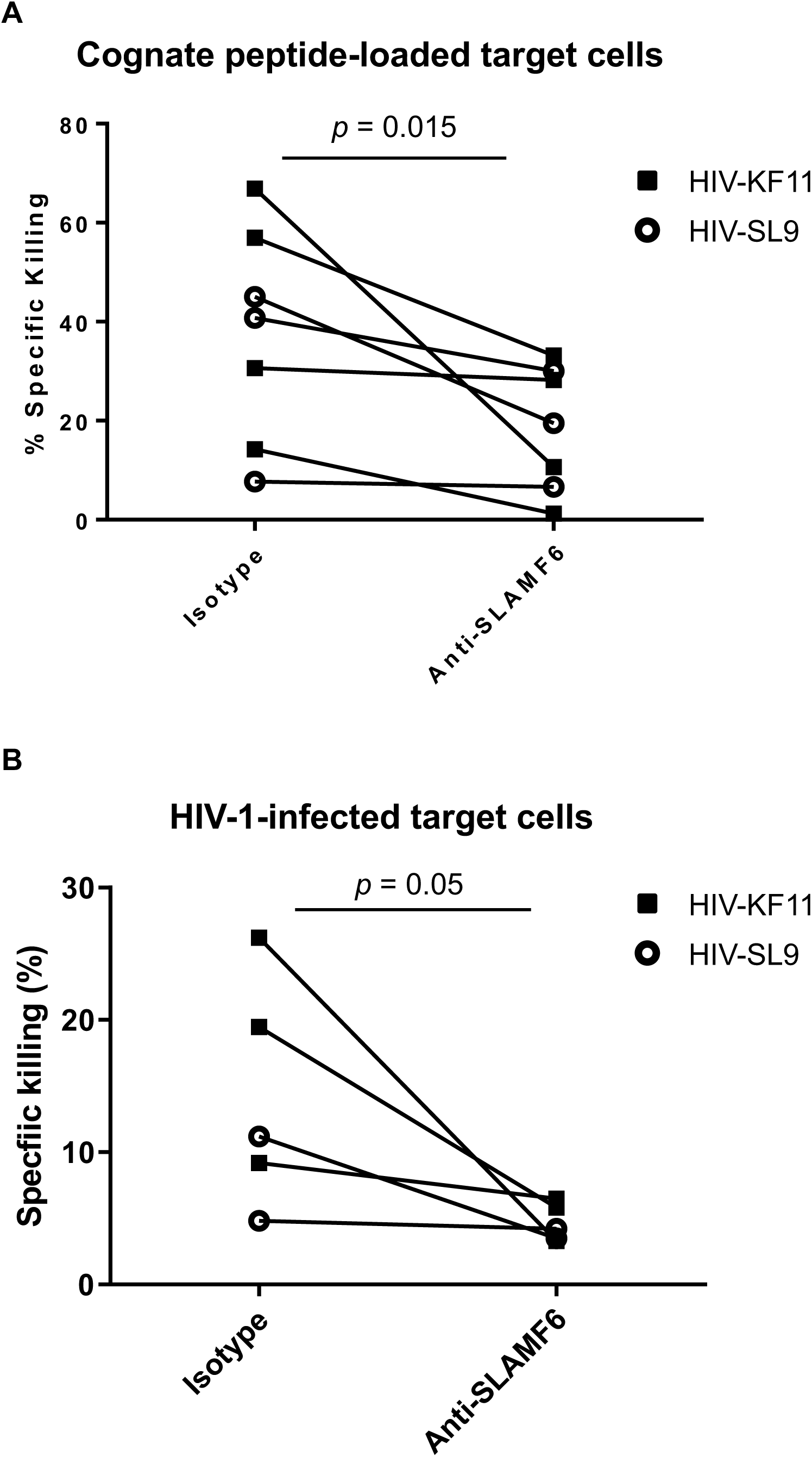
SLAMF6 is necessary for CD8^+^ T cell killing function. (**A**) Cr ^51^-labeled CD4^+^ T cells were loaded with two different peptides, either HIV-KF11 (black squares) or HIV-SL9 (open circles), at 5 μg/mL and co-cultured with autologous peptide-specific CD8^+^ T cells. Cr ^51^ release from effectors was assessed in the presence of a SLAMF6 blocking antibody or an isotype control. Seven different CTL lines were tested. (**B**) Activated primary CD4^+^ T cells were infected with HIV-1 (clone NL4-3) virus for 48 hours, then labeled with Cr ^51^ and incubated with autologous peptide-specific CD8^+^ T effector cells for 4 hours at a ratio of 1:1 in the presence of a SLAMF6 blockade or an isotype control. Five different CTL lines were tested. Black squares represent the HIV-KF11-specific CTL lines, and open circles represent the HIV-SL9-specific CTL lines tested. Specific killing of target cells in the Cr ^51^ release assay was calculated as 100x (sample release – spontaneous release) / (maximum release - spontaneous release). Statistical differences were measured using the Wilcoxon matched-pairs test with *p*-values shown on graph.

## DISCUSSION

While the pivotal role of SLAMF6 in stabilizing immunological synapses and regulating anti-viral responses has been demonstrated in mice (19, 32), its involvement in human antiviral immune responses has been understudied. Here we delved into the specific role of SLAMF6 in the cytotoxic response against CD4^+^ T cells presenting HIV-1 and CMV-derived viral peptides, as well as HIV-1-infected CD4^+^ T cells.

In line with the previous studies in mice, our findings underscore the importance of SLAMF6 for the efficient attachment of CTLs to CD4^+^ T cell targets. Blocking SLAMF6 with anti-SLAMF6 antibodies led to a significant reduction in conjugate formation. Interestingly, CTLs recognizing HIV-1 Gag epitopes restricted by HLA-B57:01 exhibited less sensitivity to SLAMF6 blockade compared to CTLs restricted by HLA-A02:01. Given that the HLA-A*02:01-restricted CTL lines are more sensitive to blocking by SLAMF6, it was challenging to study the CD8-CD4 interaction in the context of SLAMF6 blockade by confocal microscopy as we were unable to find adequate CD8-CD4 contacts to study under the microscope. For HLA-B57:01 restricted CTL, confocal microscopy unveiled that the remaining CD8^+^-CD4^+^ T cell conjugates, after anti- SLAMF6 antibody incubation, primarily stemmed from defects in actin ring formation at the immunological synapse. Moreover, SLAMF6 blockade resulted in decreased killing of HIV-1-infected CD4^+^ T cells by HIV-1-specific CTLs restricted by both HLA alleles, underscoring the broad dependence of cytolytic function on SLAMF6.

Firstly, our study reveals the important role of SLAMF6 in the formation of antigen-specific conjugates between antiviral CD8^+^ T cells and CD4^+^ T cell targets. It is consistent with previous research highlighting SLAMF6’s significance in cell-cell interactions. For instance, in mice, the absence of intracellular SLAM-associated protein (SAP) disrupts contacts between CD4^+^ T helper cells and cognate B cells, resulting in defective germinal center maintenance and reduced B cell stimulation (33–35). Notably, deletion of SLAMF6 in CD4^+^ T cells in a mouse model reverses germinal center formation defects in SAP knockout mice, demonstrating the role of SLAMF6 engagement in SAP-mediated T cell-B cell adhesion (32). Furthermore, Dragovich et al. illustrated that SLAMF6 clustering is essential for the formation of a mature immunological synapse between T cells and antigen-presenting cells, with SLAMF6 engagement triggering an activating signal (21). Additionally, bispecific anti- CD3/SLAMF6 antibodies designed to promote SLAMF6 clustering with CD3 enhance T cell activation (36).

Our study extends these findings, demonstrating that SLAMF6, in humans, is not only pivotal in regulating cell contacts in the humoral immune response but also in the cytotoxic antiviral immune response by mediating CTL-target cell adhesion in the context of lymphotropic infections like HIV-1.

Regarding conjugate formation, we observed differential sensitivity to SLAMF6 blockade between A02:01-restricted CTL lines and B57:01-restricted ones. Of particular interest in HIV-1 infection is HLA-B57:01, strongly associated with spontaneous control (37–40). In our study, we tested an HIV-peptide presented in the HLA-B57:01 context (KF11), known for its immune protective role (37)and high-avidity class-I restricted TCRs (41). High-avidity TCRs, as suggested by previous studies, may be less reliant on accessory molecules in the immunological synapse compared to low-avidity TCRs (42, 43). Thus, we propose that, for SLAMF6-expressing lymphocyte targets, low-avidity CD8^+^ T cell responses are more dependent on SLAM receptor-mediated attachment than high-avidity responses. The TCR-KF11-MHC interaction may suffice to support stable conjugation between CD8^+^-CD4^+^ T cells. However, for HLA-A02-restricted peptides tested in our study, the TCR-pMHC interaction likely lacked the strength, making CTL lines more reliant on the co-signaling molecule SLAMF6. This observation is in line with a previous finding from our laboratory (44), which demonstrated that SLAMF2-mediated modulation of the IFNγ response from CD8^+^ T cells was more pronounced in HLA-A02- restricted specificities compared to those restricted by HLA-B57. The mechanisms underlying why HLA-A02-restricted T cells are more sensitive to SLAMF6 blockade than HLA-B57-expressing cells require further investigation. It is worth noting that, in our study, HLA-B57:01-restricted CTLs, although less sensitive to SLAMF6 in terms of attachment, still depended on SLAMF6 for both immunological synapse formation and cytolytic activity. Our results suggest that HLA-B*57:01-restricted cells might establish effector-target cell contacts differently, with the TCR-pMHC interaction being dominant, while SLAM receptors fine-tune the CTL response to lymphocyte targets. However, further experiments with various peptides presented by the same HLA alleles are needed to explore the relationship between TCR-restricted avidity, synapse formation, and sensitivity to SLAMF6. Furthermore, testing different T cell clones with the same peptide-MHC specificity but with different avidities to SLAMF6 blockade would be helpful in answering this question. It is also important to consider that SLAMF6 may have a role in immune responses among elite controllers, individuals who can control HIV-1 infection below detectable levels without treatment. A comparative study between elite controllers and progressors might provide insights into this hypothesis.

Our study also investigated the role of SLAMF6 in immunological synapse formation. A mature immunological synapse involves actin polarization towards the interface (45, 46) followed by the formation of an actin ring at the contact surface between effector and target cells (31), where adhesion molecules like LFA-1 and LFA-3 accumulate (6, 7, 10, 47). This facilitates contact stabilization and directional secretion of cytotoxic perforin and granzyme (48). Our study revealed that blocking the SLAMF6 pathway disrupted synaptic architecture, as observed through confocal imaging, and impaired the killing of CD4^+^ T cells presenting peptides and HIV-infected CD4^+^ T cells by specific CD8^+^ T cells. This suggests that SLAMF6 blockade leads to the formation of a “weak synapse,” preventing the secretion of cytotoxic granules into target cells, ultimately reducing specific killing by effector T cells. This observation aligns with previous research in a mouse model, which showed that structurally fragile synapses were associated with impaired killing responses, while well-organized synapses effectively eliminated target cells (49). Furthermore, a recent study demonstrated that SLAMF1 activation enhances the anti-tumoral CD8^+^ T cell response by forming a “corolla” of SLAMF1 around the immune synapse, amplifying TCR signaling. However, engagement with PD-1 abrogates SLAMF1-mediated TCR signaling amplification (50). Building upon these results, we hypothesize that the SLAMF6 regulatory pathway is essential for establishing a mature immunological synapse between peptide-pulsed CD4^+^ T cells and autologous peptide-specific CTLs. Given that both recognition and killing of infected CD4^+^ T cells are crucial for controlling HIV-1 infection (1), SLAMF6 may play a significant role in this context. Further investigations are needed to explore the interaction between SLAMF6 and inhibitory coreceptors like PD-1.

In mice, the absence of SAP signaling through SLAM family members SLAMF6 and SLAMF4 disrupted synapse formation exclusively between cytotoxic T cells and OVA-pulsed B cells upon SLAMF6 ligand engagement. SHP-1 phosphatase binds to the phosphorylated intracellular tail of SLAMF6, rendering it incapable of forming a proper actin ring (19). In our *in vitro* human cell model, SLAMF6 blockade resulted in a more pronounced phenotype than that observed in SAP knock-out mice. In addition to defects in actin ring formation, we noted a failure of actin polarization towards the contact zone. The underlying mechanism remains to be elucidated, but we hypothesize that basal interaction between SHP-1 phosphatase and actin may exist in cytotoxic effector cells. Thus, SLAMF6 signaling might be necessary to recruit SAP to the synapse and clear actin from its center, as suggested by Chu et al. (51). Their research indicated that the interaction between SHP-1 and the SLAMF6 intracellular tail is constitutive and requires SLAMF6 crosslinking to recruit SAP. In humans, further signaling studies are warranted to validate this hypothesis.

In the context of HIV-1 infection, the viral accessory protein Vpu has been shown to downmodulate SLAMF6 on the surface of HIV-1-infected CD4^+^ T cells (24, 52). Our work uncovers a novel evasion mechanism whereby viral protein-mediated downregulation (**supplemental Figure 4**) of SLAMF6 leads to fewer CD8^+^-CD4^+^ T cell contacts, reduced formation of mature immunological synapses, and ultimately, decreased killing of HIV-infected cells. Investigating the effects of Vpu-blocking drugs on the cytotoxic response against HIV-1 mediated by SLAMF6 could yield valuable insights.

In summary, our results highlight the significance of SLAMF6 signaling in effector-target cell adhesion, synapse formation, and cytolytic killing of CD4^+^ T cells presenting HIV-1 antigens to CD8^+^ T cells. This study sheds light on the critical role of SLAMF6 in combating lymphotropic infections like HIV-1 and suggests potential interventions that could enhance the effects of therapeutic vaccines and T-cell-based immunotherapies for infectious diseases. For instance, exploring the addition of a SLAMF6 cross-linking antibody as an adjuvant to activate its signaling pathway may increase the number of effector-target cell interactions and stabilize immune synapses between HIV-infected cells and HIV-1-specific CD8^+^ T cells.

## Acknowledgements

We thank Sylvie Le Gall and Annmarie McKeon for proofreading this manuscript. We thank Thomas Diefenbach for helping with Confocal Microscopy techniques. We thank volunteer blood donors for their participation in our study. This work was supported by NIH/NIAID R56AI095088 to DGK, and the Terry and Susan Ragon Foundation. Blandine Monel was supported by Howard Hughes Medical Institute and Pedro A. Lamothe was supported by the Howard Hughes Medical Institute International Student Research Fellowship.

