## Supplemental figures for "SLAMF6 enables efficient attachment, synapse formation, and killing of HIV-1-infected CD4^+^ T cells by virus-specific CD8^+^ T cells"

**Supplemental Figure 1**

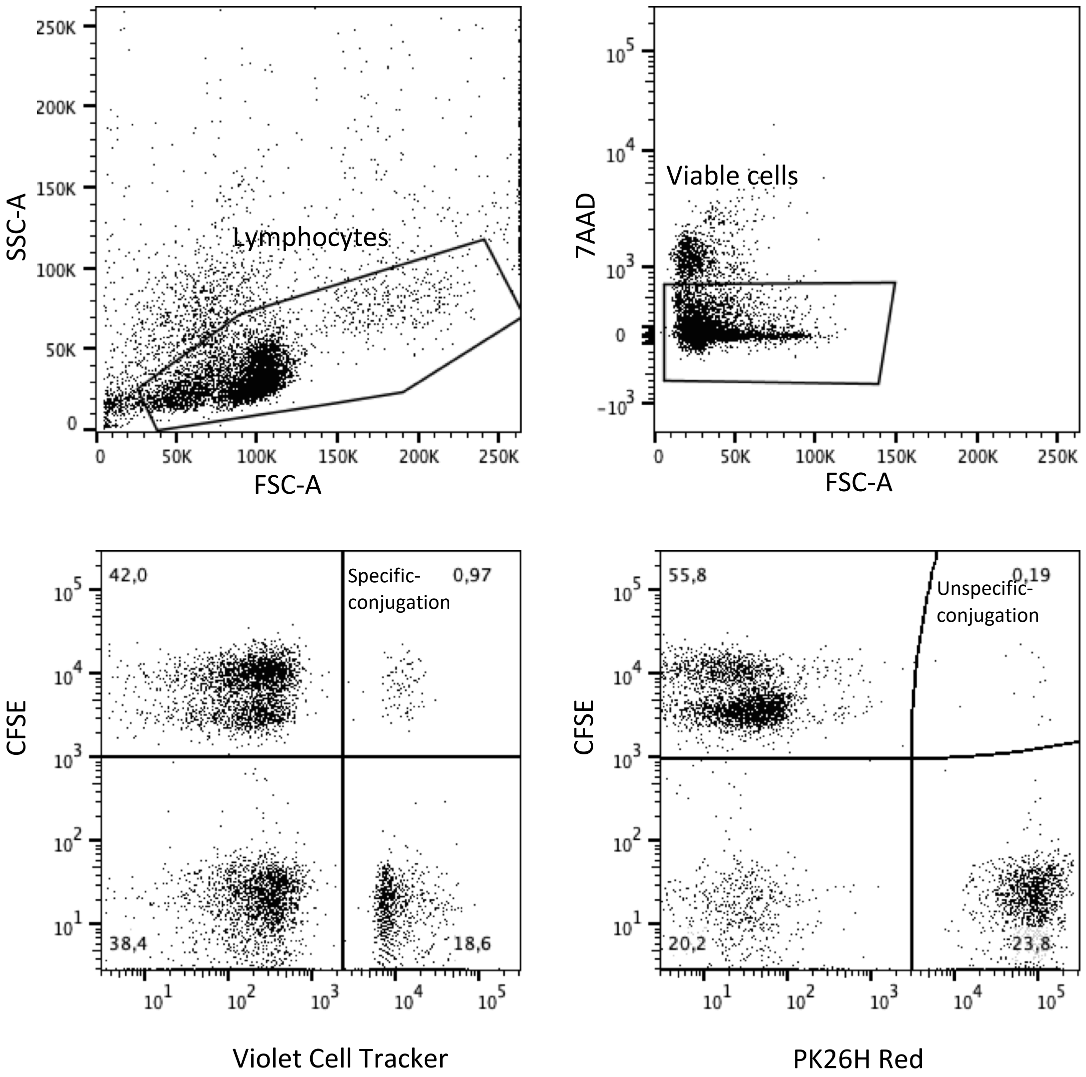


**Supplemental Figure 1: Gating strategy flow cytometry showing a representative gating strategy**. After gating on lymphocytes gate, 7AAD negative cells were selected (viable cells) and, within those the percent of specific conjugation rate ($CFSE^+$ and Violet Cell $Tracker^+$) or Unspecific conjugation rate ( $CFSE^+$ PK26H $Red^+$) was assessed.

**Supplemental Figure 2**

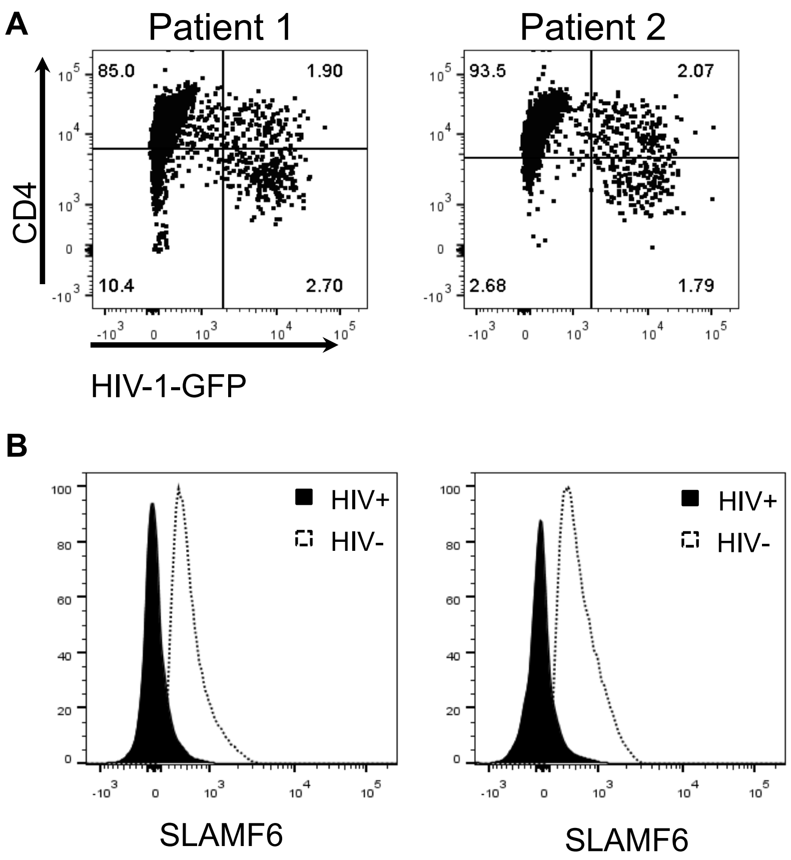


**Supplemental Figure 2:** SLAMF6 surface expression is reduced in HIV-infected CD4+ T-cells. (A) Activated primary CD4+ T-cells from two HIV-negative subjects were infected with NL4.3-GFP HIV and were stained for CD4+ T cells expression. Infected cells were gated on CD4+ T cells low and GFP+ cells. (B) Expression of SLAMF6 at the surfaces of infected GFP+cells (filled) was compared to the uninfected GFP- cells (dotted line).

### Supplemental Figure 3

**A**

| Donor ID | Age | Sex | Therapy | CD4 count (cell/μL) | Viral load (copies/mL) | Category |
|---|---|---|---|---|---|---|
| 1 | 43 | Female | None | 717 | 53,800 | Viremic (chronic progressor) |
| 2 | 45 | Male | None | 330 | 26,900 | Viremic (chronic progressor) |
| 3 | 34 | Male | None | 445 | 28,400 | Viremic (chronic progressor) |
| 4 | 44 | Female | None | 597 | 73,900 | Viremic (chronic progressor) |
| 5 | 45 | Female | None | 567 | 88,900 | Viremic (chronic progressor) |
| 6 | 51 | Female | None | 538 | 51,700 | Viremic (chronic progressor) |
| 7 | 34 | Female | None | 473 | 7,550 | Viremic (chronic progressor) |
| 8 | 46 | Male | None | 1465 | 56 | Controller (low viremia) |
| 9 | 42 | Female | HAART | 242 | <75 | HAART treated |
| 10 | 40 | Male | HAART | 713 | <20 | HAART treated |

**B**

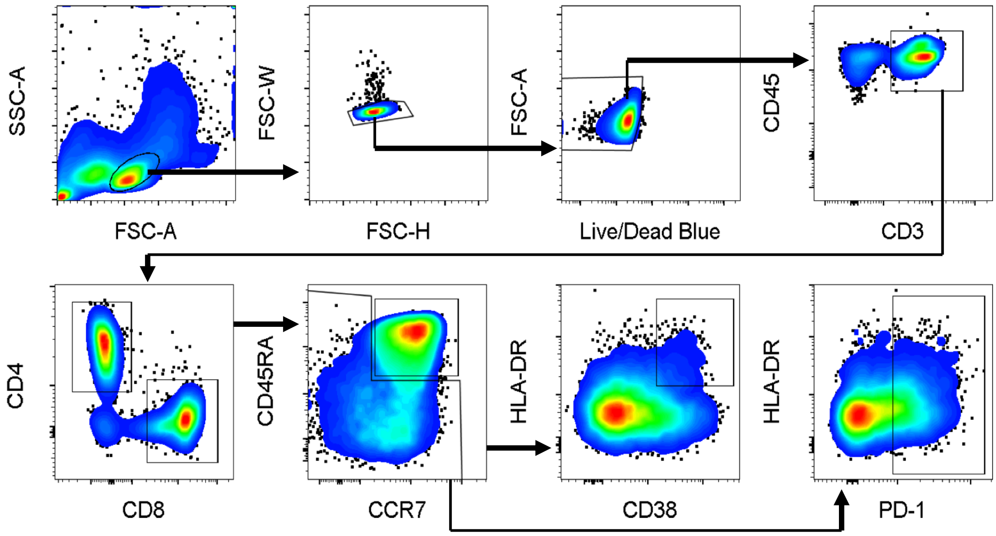


**C**

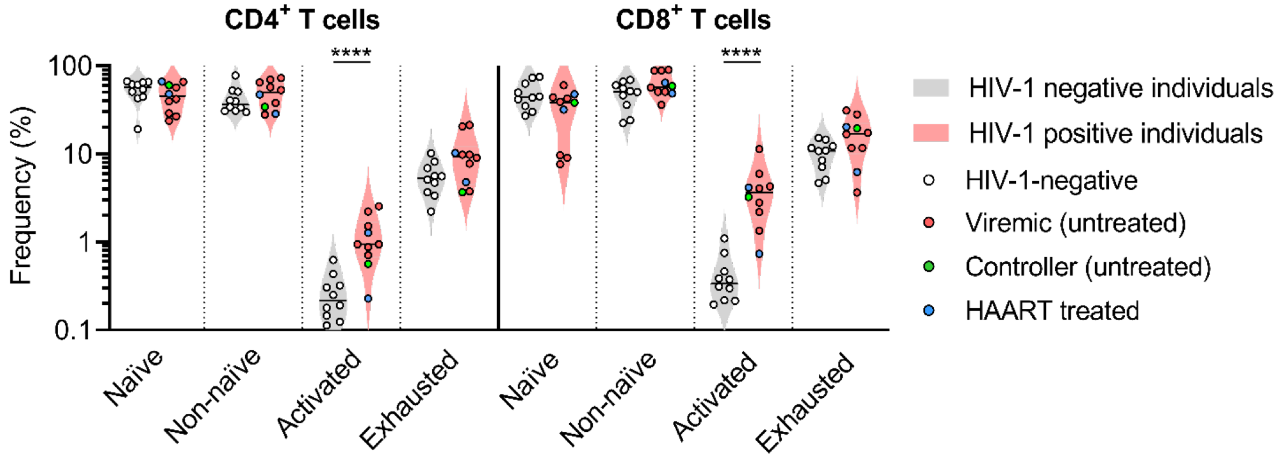


**Supplemental Figure 3**: **Characterization of SLAMF6 expression in CD4+ and CD8+ T-cell subsets among HIV-1-positive and HIV-1-negative donors.** (**A**) Demographics and relevant clinical data are shown for HIV-1-positive donors (*n* = 10); HIV-1-negative donors were healthy anonymous adults. (**B**) Flow cytometry gating scheme used to differentiate different $CD4^+$ and $CD8^+$ T-cell subsets is demonstrated. (**C**) The frequency of the different T-cell subsets among HIV-1-positive and HIV-1-negative donors is shown. One-way ANOVA with correction for multiple comparisons was performed; statistical significance is denoted as follows: **** $p < 0.0001$

**Supplemental Figure 4**

**A**

| | E:T | | | | | | Spontaneous release | | Maximum Release | |
|---|---|---|---|---|---|---|---|---|---|---|
| [KF11] | 1:1 | 1:1 | 1:4 | 1:4 | 1:8 | 1:8 | CD4 only | CD4 only | Triton | Triton |
| 5ug | 138 | 114 | 96 | 91 | 66 | 62 | 19 | 21 | 224 | 220 |
| 5ug | 95 | 84 | 87 | 102 | 58 | 80 | 27 | 25 | 261 | 247 |
| no peptide | 18 | 13 | 23 | 23 | 13 | 24 | 14 | 36 | 185 | 188 |
| no peptide | 19 | 24 | 24 | 17 | 24 | 19 | 23 | 23 | 188 | 218 |

**B**

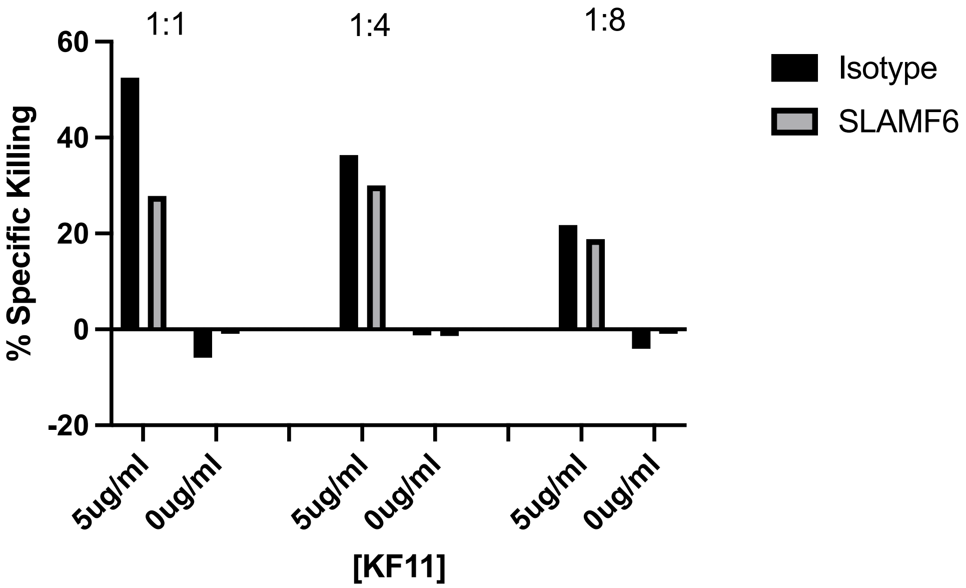


**Supplemental Figure 4: Set-up of the chromium release killing assay.** We tested various effector-to-target (E:T) cell ratios. Target cells were either loaded with 5 µg/ml of peptide or left unloaded before the addition of CTL effector cells. The percentage of killing was determined using the formula: 100 x (average of the two duplicates −average of spontaneous release)/(average duplicates of maximum release−average of spontaneous **release). (A)** Raw data obtained from a Chromium release assay. (**B)** The percentages of killing under different experimental conditions are presented below.
